# Predicting The Pathway Involvement Of Metabolites Based on Combined Metabolite and Pathway Features

**DOI:** 10.1101/2024.04.01.587582

**Authors:** Erik D. Huckvale, Hunter N.B. Moseley

**Affiliations:** Markey Cancer Center, University of Kentucky, Lexington, KY 40506, USA; Superfund Research Center, University of Kentucky, Lexington, KY 40506, USA; Department of Toxicology and Cancer Biology, University of Kentucky, Lexington, KY 40536, USA; Department of Molecular and Cellular Biochemistry, University of Kentucky, Lexington, KY 40506, USA; Institute for Biomedical Informatics, University of Kentucky, Lexington, KY 40506, USA

**Keywords:** Metabolism, Metabolite, Metabolic pathway, Machine learning, Deep learning, XGBoost, Multi-layer perceptron, Supervised learning, Binary classification

## Abstract

A major limitation of most metabolomics datasets is the sparsity of pathway annotations of detected metabolites. It is common for less than half of identified metabolites in these datasets to have known metabolic pathway involvement. Trying to address this limitation, machine learning models have been developed to predict the association of a metabolite with a “pathway category”, as defined by one of the metabolic knowledgebases like the Kyoto Encyclopedia of Gene and Genomes. Most of these models are implemented as a single binary classifier specific to a single pathway category, requiring a set of binary classifiers for generating predictions for multiple pathway categories. This single binary classifier per pathway category approach both multiplies the computational resources necessary for training while diluting the positive entries in gold standard datasets needed for training. To address the limitations of training separate classifiers, we propose a generalization of the metabolic pathway prediction problem using a single binary classifier that accepts both features representing a metabolite and features representing a generic pathway category and then predicts whether the given metabolite is involved in the corresponding pathway category. We demonstrate that this metabolite-pathway features-pair approach is not only competitive with the combined performance of training separate binary classifiers, but it outperforms the previous benchmark models.

## 1. Introduction

Metabolism is the set of biochemical processes within cells and organisms that sustain life. Metabolites are chemical compounds that take part as reactants and/or products of chemical reactions involved in metabolism. The products of one reaction can act as reactants in another, resulting in chains of reactions occurring in different parts of a cell or organism, diverging in different directions, and serving different metabolic purposes. These reaction chains have been organized into networks of connected reactions known as biochemical pathways. These pathways have been grouped into broader pathway categories, classified by the types of reactants and products involved, cellular location, metabolic purpose, etc. Since metabolites are the reactants and products of metabolic reactions and reactions are the building blocks of pathways, certain metabolites are necessarily involved in certain pathway categories. However, the entirety of metabolic pathways are not known since not all of the chemical reactions performed in cellular metabolism have been discovered. This results in a recurring problem faced by biologists, biochemists, and bioinformaticians which is coming across metabolites lacking information related to their pathway involvement.

Knowledgebases such as the Kyoto Encyclopedia of Genes and Genomes (KEGG) [1–3] and BioCyc, particularly via their database called MetaCyc [4], have made tremendous contributions to creating databases containing metabolites with pathway annotations. However, due to the costly and laborious nature of experimentally determining the pathway involvement of metabolites, many of the metabolites in such metabolic knowledge-bases are still not annotated or are only partially annotated to metabolic pathways. For example, as of July 3^rd^ 2023, a total of 19,119 compounds existed in the KEGG database with only 6,736 of them having annotated pathway involvement [5]. Because of the lack of annotation, several machine learning methods have been proposed to predict the pathway categories that a metabolite is involved in given information about the metabolite’s chemical structure. There have been several publications in particular that have trained supervised learning models, most notably graph neural networks [6], on compounds in the KEGG database. Models were trained and evaluated on datasets with metabolites as entries, the features being chemical structure information of these metabolites and the labels being high level metabolic pathway categories. Specifically, KEGG provides a hierarchy of broader pathway categories branching into more granular pathway categories as seen here: https://www.genome.jp/brite/br08901. There are 12 categories under metabolism, namely 1. Amino acid metabolism 2. Biosynthesis of other secondary metabolites, 3. Carbohydrate metabolism, 4. Chemical structure transformation maps, 5. Energy metabolism, 6. Glycan biosynthesis and metabolism, 7. Lipid metabolism, 8. Metabolism of co-factors and vitamins, 9. Metabolism of other amino acids, 10. Metabolism of terpenoids and polyketides, 11. Nucleotide metabolism, and 12. Xenobiotics biodegradation and metabolism. Several past publications trained models to predict 11 out of the 12 categories, excluding ‘Chemical structure transformation maps’, likely due to its difficulty to predict. Since some of these publications were proven to be invalid, using a dataset that contained duplicate entries in both the train and test set [7], Huckvale et al developed a new KEGG-derived benchmark dataset for the task of developing models for predicting pathway category involvement for all 12 of the aforementioned categories based on metabolite chemical structure [5]. Huckvale et al also demonstrated a set of binary classification models trained on this benchmark dataset for predicting pathway category involvement.

However, these benchmark models involved training a separate model for each pathway category. This approach complicates the design, implementation, and maintenance of the model training, evaluation, and deployment pipeline. The amount of computational resources required are multiplied by the number of models needed, which is one for each pathway category. These shortcomings are exacerbated further when proposing to train models on more granularly-defined pathway categories as compared to the most highlevel pathway categories. For example, KEGG defines 12 topmost pathway categories, but defines 184 in total when descending one level down the hierarchy. The current benchmark dataset includes 5,683 metabolites with pathway annotations and chemical structure representations, which is adequate to train 12 pathway category-specific models, but is woefully inadequate to train 184 separate models, due to the diminishing small number of positive entries for each pathway. Additionally, the resulting models are only equipped to predict on the specific pathway category they were trained on. There are several other databases and use cases that may not organize pathway category in the exact same way as the hierarchy found in KEGG. For example, we see that MetaCyc provides an entirely different pathway hierarchy as seen here: https://metacyc.org/META/class-tree?ob-ject=Pathways. This suggests the need for a more generic model.

In this work, we present a single binary classifier for predicting the pathway involvement of metabolites. The dataset provided by the work of Huckvale et al [5] uses an atom coloring technique [8] to represent the sub-structures of molecules, the features of a metabolite being the count of such molecular substructures that are present in the molecule. Every pathway category has certain metabolites associated with it, so a pathway category can be generically represented by aggregations of the features of the metabolites associated with the pathway category. Given chemical structure information of metabolites along with structured informational representations of pathway categories, we have constructed a dataset where entries consist of metabolite features paired with pathway features. With a dataset of these metabolite-pathway feature-pairs, we have trained and evaluated models to predict whether the metabolite of the pair is involved in the pathway category of the pair. This technique uses just one model that can effectively predict the original 12 pathway categories and paves the way for a generic classifier that can predict the presence of metabolites in arbitrary pathway categories. We demonstrate that not only does the metabolite-pathway-pairs technique perform well compared to training a separate model per classifier, our best model has even exceeded the combined performance of the prior benchmark models, not just in average MCC, but with a significantly lower standard deviation.

## 2. Materials and Methods

### 2.1. Generating the feature vectors

Huckvale et al [5] previously generated a dataset of 5,683 entries, each entry containing a vector of atom-color [8] features corresponding to a metabolite. Building off of this dataset of metabolite features, we constructed pathway features via the process in Figure 1. Each of the 12 pathway categories have a subset of the 5,683 available metabolites that are associated with it (e.g. ‘Amino acid metabolism’ contains 611 out of the 5,683 metabolites, ‘Biosynthesis of other secondary metabolites’ contains a different subset of 1,486 metabolites, etc.). With each subset, we created vectors that are sums of the features of the metabolites associated with the corresponding pathway category. While the features of a single metabolite are the counts of the atom colors within it, the resulting summed features are the number of occurrences of the atom colors across all the metabolites within the entire pathway category. However, we could not simply use these raw counts for the pathway features since different pathway categories have different amounts of metabolites within them. To correct this, we applied a bond inclusivity-specific soft max normalization by calculating the proportion of each atom color within each pathway category and dividing the raw counts by the total count of the other atom colors within the same pathway category and of the same level of bond inclusion (Figure 1). For example, for the elemental atom colors (0-bond-inclusion), if the total count of elemental atom colors for a given pathway category was 10,000 but the count for the carbon atom was 1,000, then the feature value for the carbon atom color in that pathway category would be 0.1. This effectively normalizes the pathway categories such that they become comparable to each other despite one category having more metabolites than another. For consistency, we also normalized the metabolite features in the same way. The resulting metabolite features were the proportion of occurrences of each atom color compared to every other atom color of the same bond inclusion level within a single compound and the resulting pathway features were likewise the proportion of occurrences of that atom color across compounds within the entire pathway category (Figure 1).

**Figure 1.**
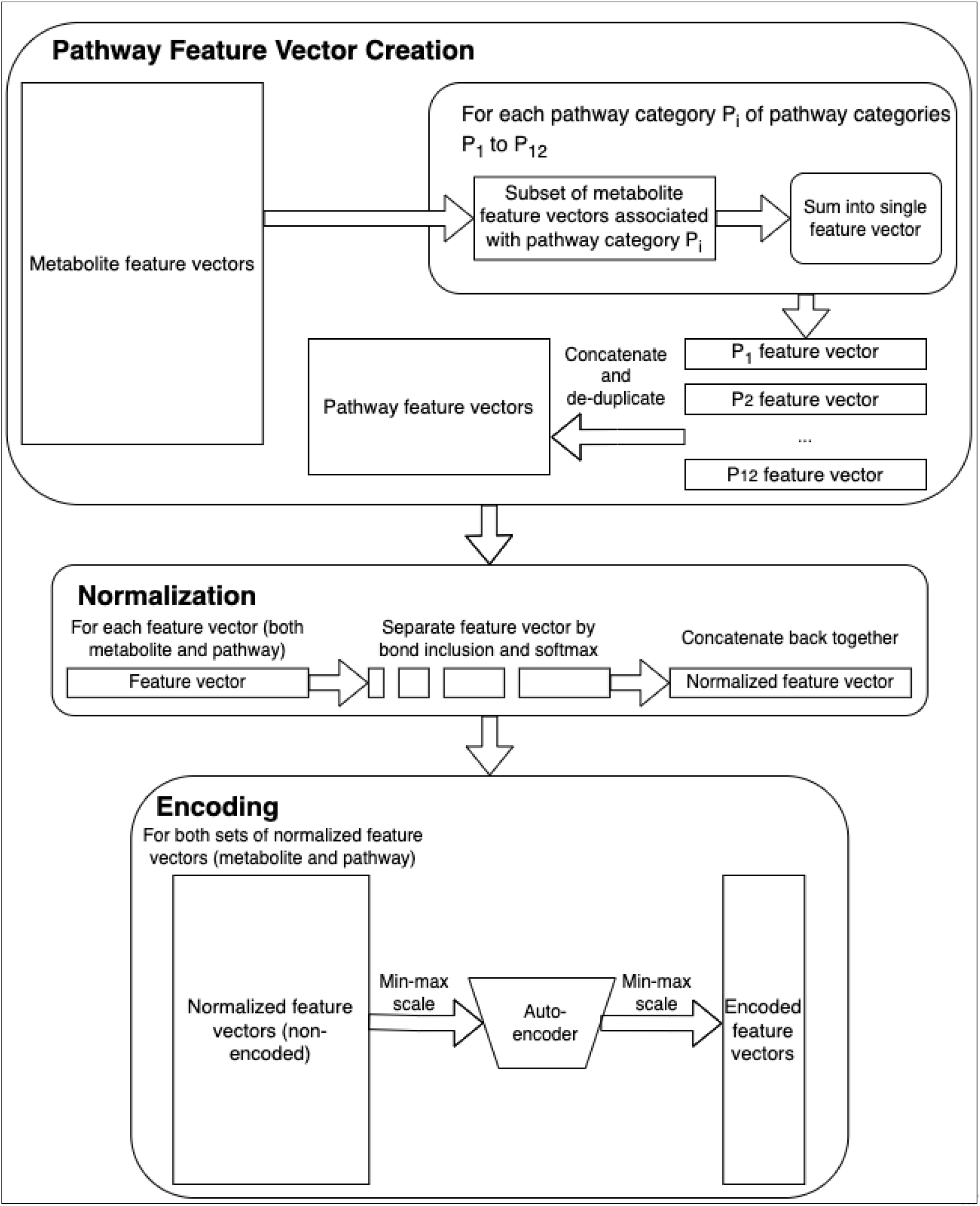
Feature engineering.

Finally, upon creating the 12 pathway feature vectors, there were duplicate pathway features across the 12 vectors. We created a copy of the set of the pathway features with the duplicate features removed. The original set (containing duplicate features) was used to make encoded features using an auto-encoder (Figure 1) while the de-duplicated set was used to train models directly. While auto-encoders [9] have been known to improve classification performance via feature reduction, particularly reducing redundant features, they at the very least reduce training time and other computational resources. We created encoded counterparts of the metabolite and pathway feature sets to determine if our models would perform at least as well after passing through an auto-encoder. While we normalized entry-wise initially, we additionally normalized feature-wise prior to training the auto-encoder via min-max scaling (Figure 1). The auto-encoder was trained on both the metabolite and pathway features since they had the same atom colors (the original set of pathway features were used by the auto-encoder while the de-duplicated set of pathway features were used downstream). We performed min-max scaling again after passing the non-encoded data through the auto-encoder to create the final encoded features (Figure 1).

Table 1 provides characteristics of the individual metabolite feature sets and pathway feature sets before they were paired together. Since we built off of the work of Huckvale et al [5], the number of metabolite entries and features were the same as with their work, the metabolite features having already been de-duplicated. Since the pathway features were derived from the metabolite features, the number of pathway features was initially the same as that of the metabolite features. De-duplicating the pathway features removed 9,220 duplicate features from the original 14,655. Such a large proportion of the features were duplicates likely because we were considering just 12 pathway categories as compared to 5,683 metabolites. Since the pathway features were not de-duplicated before passing through the auto-encoder, the autoencoder could encode both the pathway features and metabolite features (auto-encoders expect their input to always be the same size). Encoding the features to one tenth their original size resulted in both sets having 1,465 encoded features (Table 1).

**Table 1.** Characteristics of the individual feature sets.

| Feature type | Encoded | Number of entries | Number of features |
| --- | --- | --- | --- |
| Metabolite | No | 5,683 | 14,655 |
| Metabolite | Yes | 5,683 | 1,465 |
| Pathway | No | 12 | 5,435 |
| Pathway | Yes | 12 | 1,465 |

### 2.2. Constructing the metabolite-pathway feature-pair dataset via cross-join

Once all four feature sets were complete, we fed the data to the machine learning models as a cross join between the metabolite feature vectors and the pathway feature vectors (i.e. every metabolite feature vector was joined with every pathway feature vector). Concatenating the two vectors together resulted in sets of metabolite-pathway feature-pair entries (Figure 2). The corresponding label of each entry was a binary label indicating whether the given metabolite was part of the given pathway category (positive entry) or not (negative entry).

**Figure 2.**
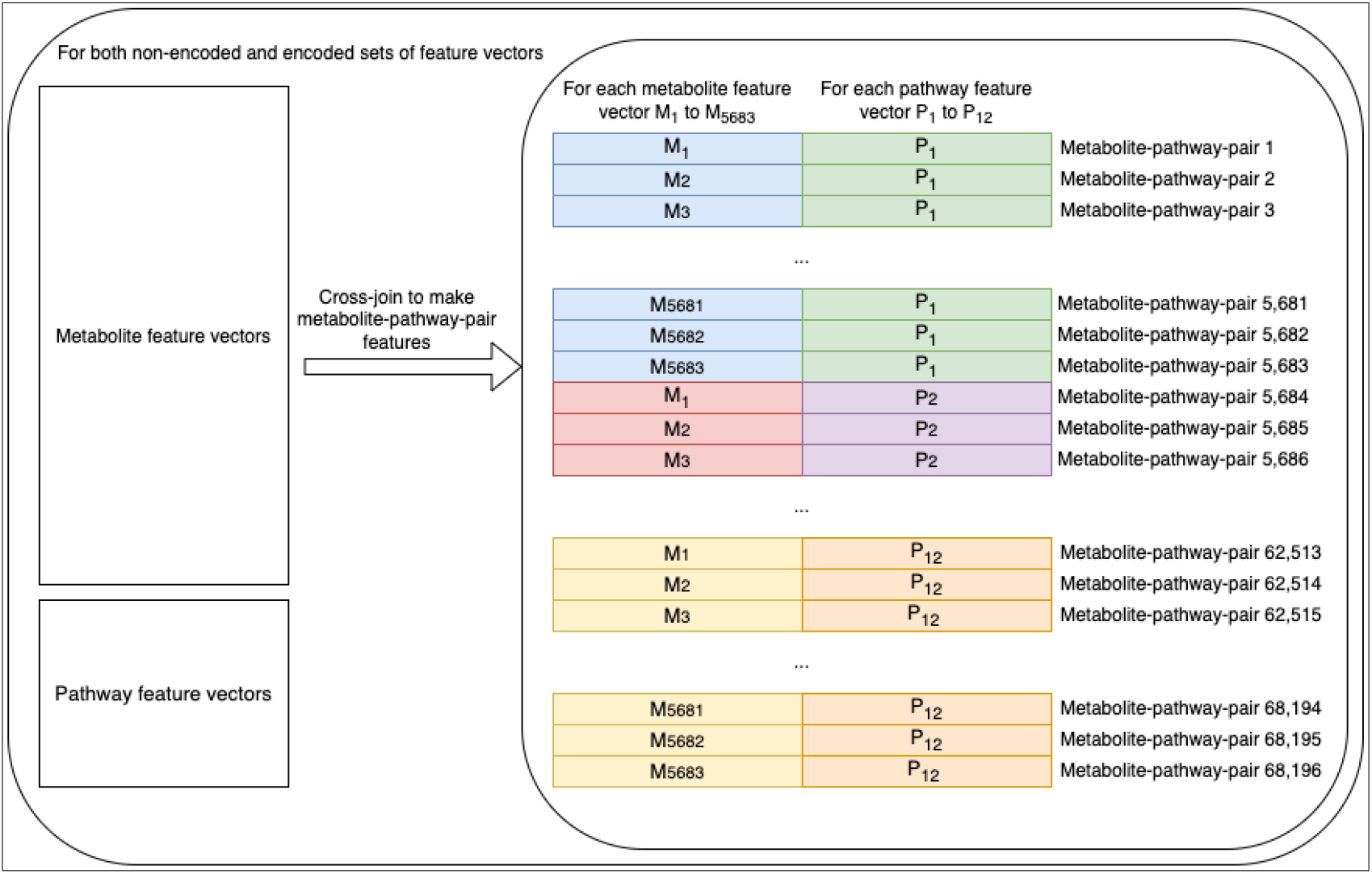
Creation of the metabolite-pathway feature-pair sets via cross-join.

Table 2 shows that the characteristics of the non-encoded metabolite-pathway feature-pair dataset were all the same as the encoded counterpart except for the number of features, which of course, were one tenth the amount as the non-encoded. The resulting number of entries was the number of metabolite entries (5,683) multiplied by the number of pathway entries (12). The number of positive entries was the sum of the number of metabolites associated with each of the 12 pathway categories. While each pathway category had a different number of metabolites associated, the combination of them all resulted in an overall proportion of about 10.6% positive entries (Table 2).

**Table 2.**
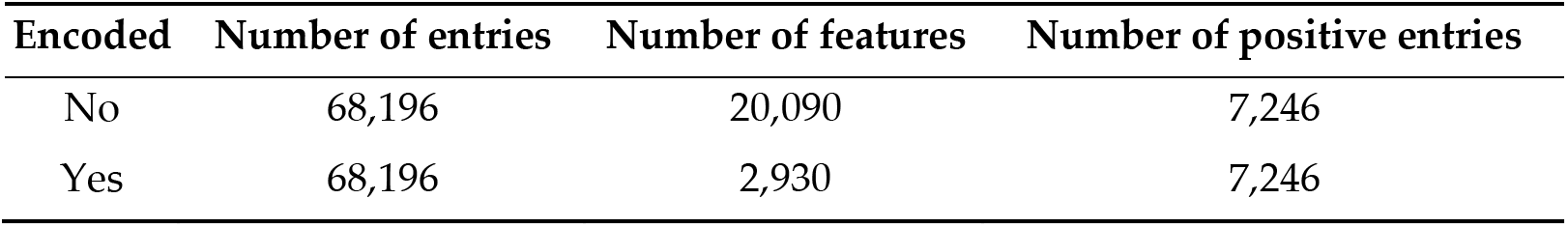
Characteristics of the metabolite-pathway feature-pair datasets after the cross-join.

### 2.3. Hyperparameter tuning and model evaluation

While the XGBoost [10] model previously performed best overall in the work of Huckvale et al [11], we suspected that a neural network approach may perform better than before considering the increase in data as a result of the cross join (68,196 entries compared to 5,683). Therefore, we ran experiments using both an XGBoost model as well as a multi-layer perceptron (MLP) [12]. Figure 3 shows the overview of hyperparameter tuning, model training, and model evaluation using both the non-encoded and encoded features. This included performing 100 trials of hyperparameter tuning for each of the four combinations of model and feature set, using the Optuna Python library [13]. For each trial, we performed up to 20 cross-validation (CV) iterations (some trials had less than 20 if they were pruned due to not showing promise), creating a stratified train-test split [14] for each iteration. With a fairly low proportion of positive entries (Table 2), we decided to duplicate the positive entries in the training sets until the proportion of positive entries was equal or just under 50%. The test sets, however, retained the same proportion of positive entries, since duplicate entries in a test set can lead to overly optimistic and otherwise misleading results. Upon training the model, we obtained predictions on the test set and compared them against the labels of the same, calculating the Matthew’s correlation coefficient (MCC) [15,16] of each train-test split. The median MCC across the CV iterations was used to determine the most successful hyperparameter tuning trial. Table S1 shows the hyperparameters selected from the best trial for each model-feature-set combination.

**Figure 3.**
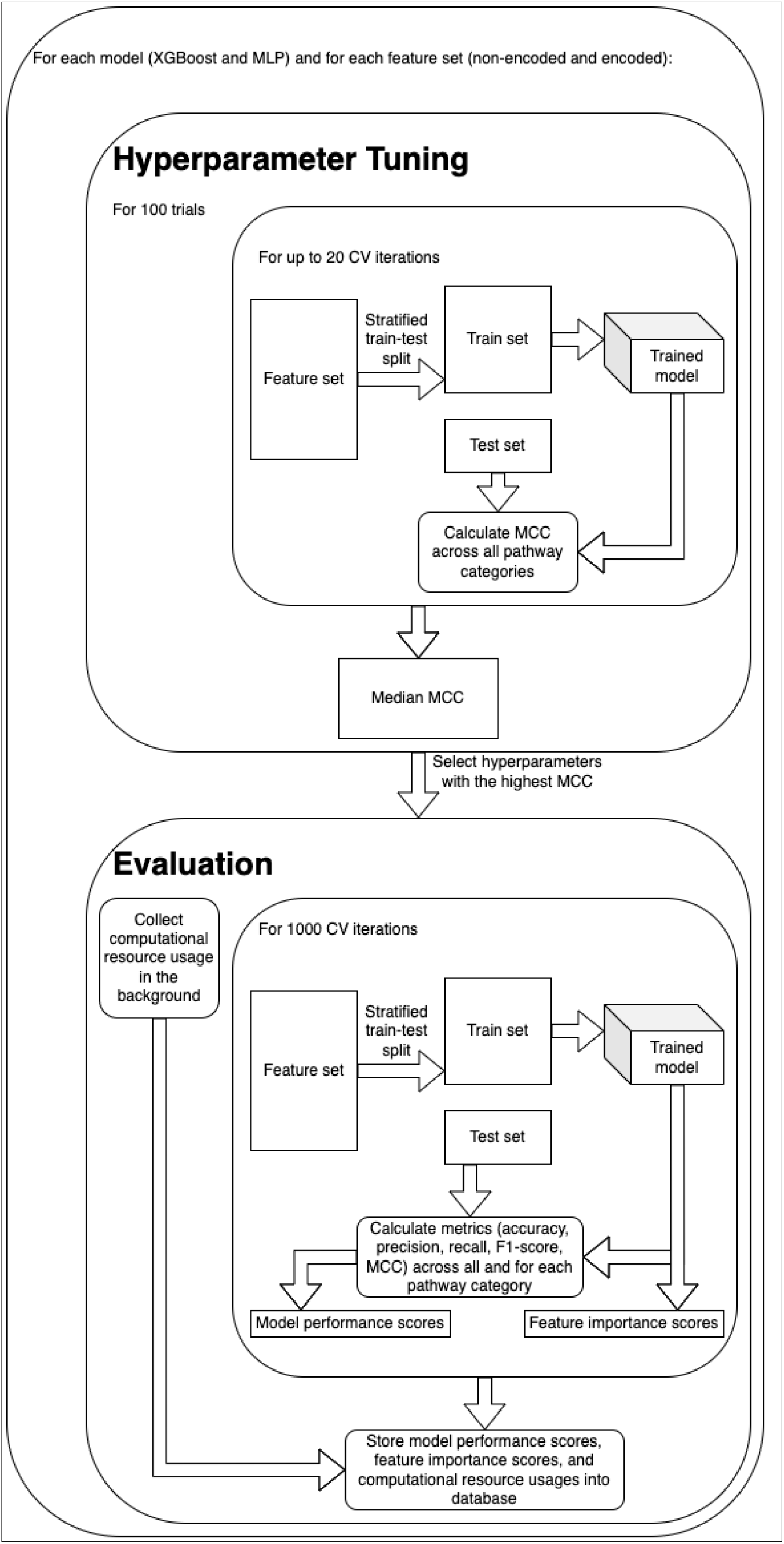
Overview of hyperparameter tuning, model training, and model evaluation.

Using the best hyperparameters for each model, we trained the models over 1,000 CV iterations, performing train-test splits similar to those made during hyperparameter tuning but performing 1,000 iterations instead of 20 for the final evaluation. Pragmatically, we only used 20 for hyperparameter tuning to save time, expecting that 20 would provide a reasonable estimate of overall model performance. We calculated five model performance metrics on the test set for each CV iteration including accuracy, precision, recall, F1-score, and MCC. Additionally, we measured the importance of each input feature, but only for XGBoost trained on the non-encoded feature set. We stored the model performance scores and the feature importance scores in a database file to summarize and visualize the results downstream (Figure 3).

For tuning the hyperparameters and the final CV analyses of both the XGBoost and MLP, weused high performance computing (HPC) machines with a system capacity of 187GB of RAM and 32 cores per node, the CPUs being ‘Intel® Xeon® Gold 6130 CPU@2.10GHz’. Using the SLURM HPC job manager, no more than 72 hours of compute time was allocated for each of the four hyperparameter tunings and CV analyses. The XGBoost runs allocated 10 cores with 17 gigabytes of RAM allocated per core. The MLP runs allocated 16 cores with 6 gigabytes of RAM per core. Both of the XGBoost runs used a GPU with 12GB of GPU memory, thenameofthe GPU card being‘Tesla P100 PCIe 12GB’.

All scripts for data processing and analysis were written in the Python programming language [17] and results were stored in an SQL [18] database managed by the DuckDB Python package [19]. Summarization and visualization of said results were performed using the Tableau business intelligence software [20] and the Seaborn [21] Python package (built on the MatPlotLib [22] python package) within Jupyter notebooks [23]. Data processing was facilitated by the NumPy [24], Pandas [25], and H5Py [26] Python packages. Evaluation metrics were computed using the Sci-Kit Learn [27] Python package. Pearson and Spearman correlation coefficients were computed using the SciPy [28] Python package. The XGBoost model was implemented using the XGBoost Python package [10], while the auto-encoder and MLP were implemented using the Pytorch Lightning [29] and Torch Geometric [30] packages built on top of the PyTorch [31] package. Computational resource profiling was performed using the gpu-tracker Python package (***<u>CITE GPU-TRACKER</u> <u>PAPER</u>***).

## 3. Results

### 3.1. Model Performance

Table 3 provides the average and standard deviation of MCC for each combination of model (MLP and XGBoost) and feature set (encoded by the auto-encoder or not) and for each of the 12 pathway categories. See Table S2 for all metrics in addition to MCC.

**Table 3.** Model performance by pathway category.

| Model | Pathway Category | Average MCC | Standard Deviation |
| --- | --- | --- | --- |
| MLP | Amino acid metabolism | 0.6712 | 0.0518 |
|  | Biosynthesis of other secondary metabolites | 0.8268 | 0.0268 |
|  | Carbohydrate metabolism | 0.7736 | 0.0504 |
|  | Chemical structure transformation maps | 0.3879 | 0.0696 |
|  | Energy metabolism | 0.5707 | 0.1035 |
|  | Glycan biosynthesis and metabolism | 0.7560 | 0.0589 |
|  | Lipid metabolism | 0.9029 | 0.0320 |
|  | Metabolism of cofactors and vitamins | 0.7403 | 0.0481 |
|  | Metabolism of other amino acids | 0.5937 | 0.0813 |
|  | Metabolism of terpenoids and polyketides | 0.8867 | 0.0251 |
|  | Nucleotide metabolism | 0.7680 | 0.0816 |
|  | Xenobiotics biodegradation and metabolism | 0.8677 | 0.0305 |
| MLP<br>with<br>Encoding | Amino acid metabolism | 0.6327 | 0.0520 |
|  | Biosynthesis of other secondary metabolites | 0.8110 | 0.0296 |
|  | Carbohydrate metabolism | 0.7694 | 0.0467 |
|  | Chemical structure transformation maps | 0.4789 | 0.0621 |
|  | Energy metabolism | 0.5814 | 0.0982 |
|  | Glycan biosynthesis and metabolism | 0.7650 | 0.0565 |
|  | Lipid metabolism | 0.8930 | 0.0292 |
|  | Metabolism of cofactors and vitamins | 0.7001 | 0.0517 |
|  | Metabolism of other amino acids | 0.5733 | 0.0812 |
|  | Metabolism of terpenoids and polyketides | 0.8741 | 0.0275 |
|  | Nucleotide metabolism | 0.7571 | 0.0856 |
|  | Xenobiotics biodegradation and metabolism | 0.8339 | 0.0347 |
| XGBoost | Amino acid metabolism | 0.6110 | 0.0525 |
|  | Biosynthesis of other secondary metabolites | 0.7803 | 0.0293 |
|  | Carbohydrate metabolism | 0.7857 | 0.0454 |
|  | Chemical structure transformation maps | 0.4221 | 0.0722 |
|  | Energy metabolism | 0.5578 | 0.1090 |
|  | Glycan biosynthesis and metabolism | 0.7701 | 0.0599 |
|  | Lipid metabolism | 0.8938 | 0.0276 |
|  | Metabolism of cofactors and vitamins | 0.7157 | 0.0491 |
|  | Metabolism of other amino acids | 0.5042 | 0.0856 |
|  | Metabolism of terpenoids and polyketides | 0.8628 | 0.0271 |
|  | Nucleotide metabolism | 0.7729 | 0.0807 |
|  | Xenobiotics biodegradation and metabolism | 0.8226 | 0.0309 |
| XGBoost<br>with<br>Encoding | Amino acid metabolism | 0.4342 | 0.0628 |
|  | Biosynthesis of other secondary metabolites | 0.6897 | 0.0344 |
|  | Carbohydrate metabolism | 0.6723 | 0.0542 |
|  | Chemical structure transformation maps | 0.2580 | 0.0803 |
|  | Energy metabolism | 0.4312 | 0.1211 |
|  | Glycan biosynthesis and metabolism | 0.7040 | 0.0653 |
|  | Lipid metabolism | 0.8045 | 0.0387 |
|  | Metabolism of cofactors and vitamins | 0.5504 | 0.0594 |
|  | Metabolism of other amino acids | 0.3589 | 0.0990 |
|  | Metabolism of terpenoids and polyketides | 0.7717 | 0.0338 |
|  | Nucleotide metabolism | 0.5465 | 0.1184 |
|  | Xenobiotics biodegradation and metabolism | 0.6800 | 0.0428 |

Figure 4 provides a violin plot of the MCC obtained over the 1,000 CV iterations for the MLP models for each pathway category. The distribution of performance of the model trained on the non-encoded feature set is shown side by side with that of the corresponding model trained on the encoded feature set. We see that for most pathway categories, the model trained on the non-encoded set outperformed the encoded counterpart. However, for a few pathway categories, namely ‘Chemical structure transformation maps’, ‘Glycan biosynthesis and metabolism’, and ‘Energy metabolism’, the MLP trained on the encoded set performed better.

**Figure 4.**
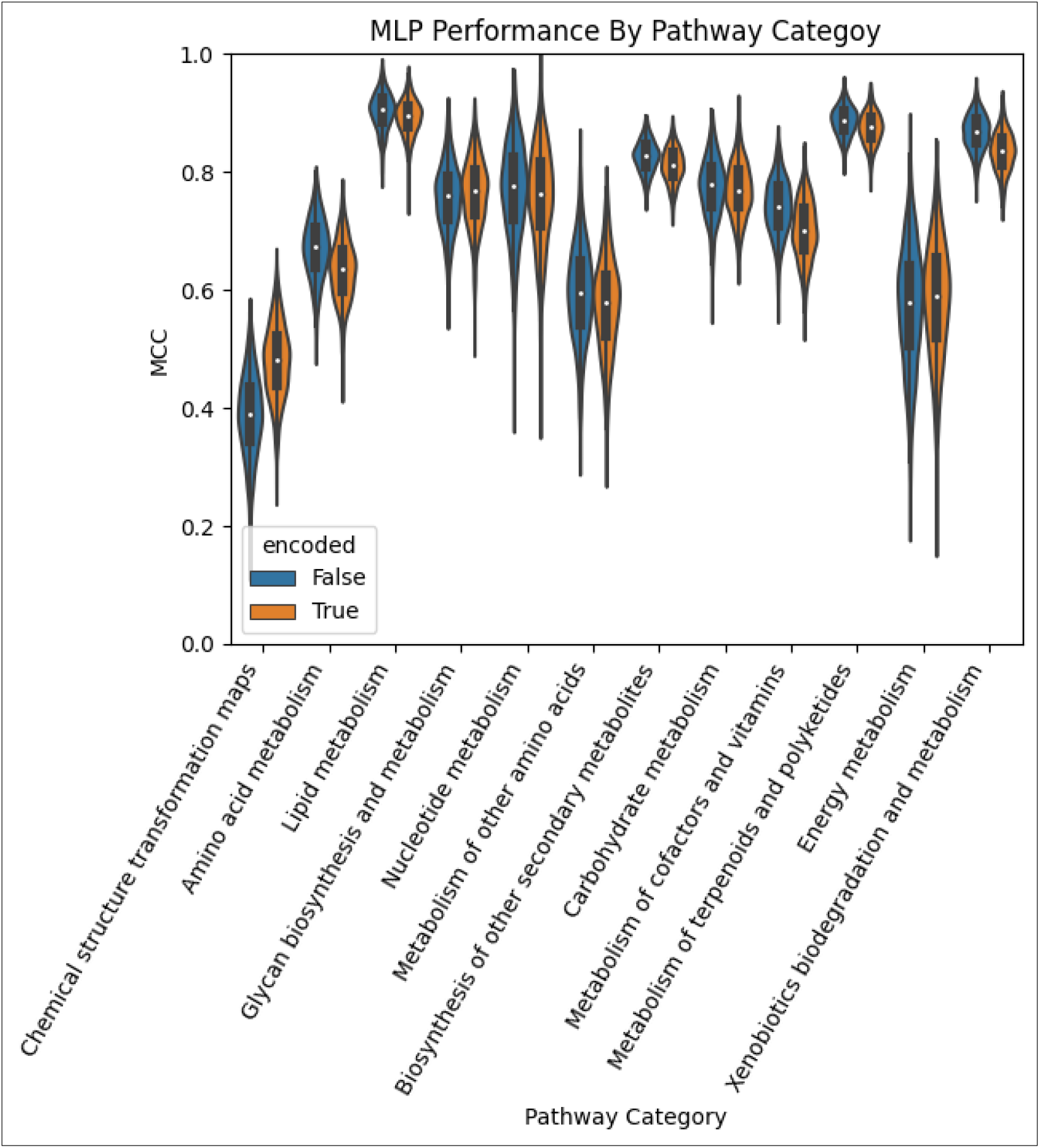
Performance of the MLP models by pathway category.

Figure 5 provides the same as Figure 4 but for the XGBoost models. We see that XGBoost consistently performs significantly worse when trained on the encoded feature set.

**Figure 5.**
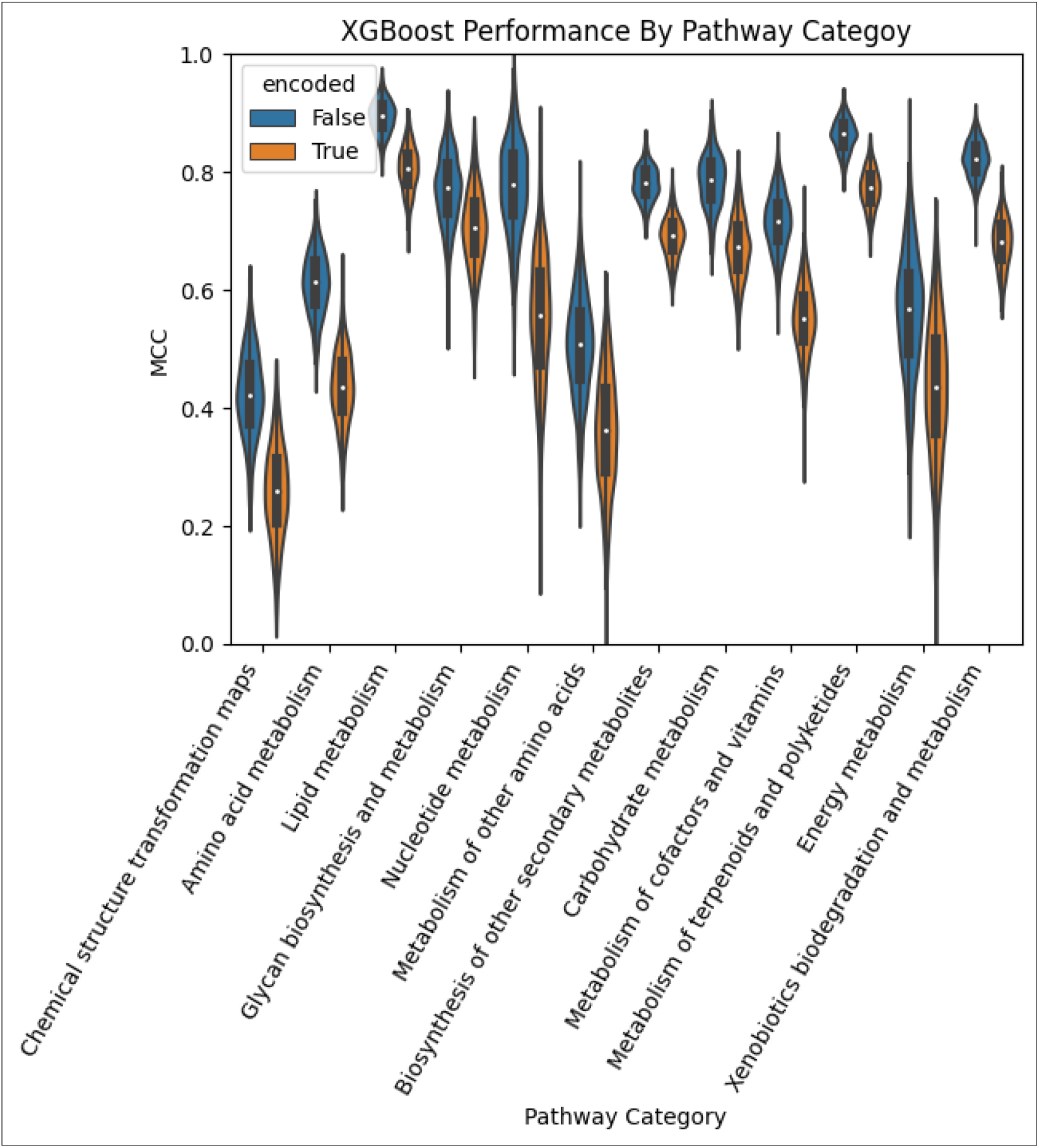
Performance of the XGBoost models by pathway category.

Table 4 compares the average and standard deviation MCC of each model and feature set combination. When compared to the previous work of Huckvale et al [5], which trained a separate binary classifier for each pathway category, we see that the MLP greatly improved when training a single binary classifier on metabolite-pathway feature-pairs. The MLP trained on the encoded data performed comparably well to the XGBoost model trained on non-encoded data which likewise performed comparably well to the XGBoost of the previous work. The XGBoost trained on the encoded data performed significantly worse than all other models and is included for completeness. While only the MLP trained on the non-encoded feature set significantly exceeded the average MCC of the best model in the previous work, the standard deviation of all metabolite-pathway-pair models are lower by an order of magnitude from those of the previous work.

**Table 4.** Performance by model.

| Model | Average MCC | Standard Deviation | Weighted Average MCC of Huckvale et al [5] | Weighted Standard Deviation |
| --- | --- | --- | --- | --- |
| MLP | 0.7844 | 0.0129 | N/A | N/A |
| MLP with Encoding | 0.7695 | 0.0139 | 0.7240 | 0.1615 |
| XGBoost | 0.7637 | 0.0126 | 0.7677 | 0.1540 |
| XGBoost with Encoding | 0.6567 | 0.0152 | N/A | N/A |

Figure 6 provides violin plots showing the distribution of the models across all pathway categories. We see that the distributions for the XGBoost trained on the non-encoded data does not even overlap with that trained on the encoded data. For the MLP, the two distributions do overlap but the MLP trained on the non-encoded data is clearly the highest performing.

**Figure 6.**
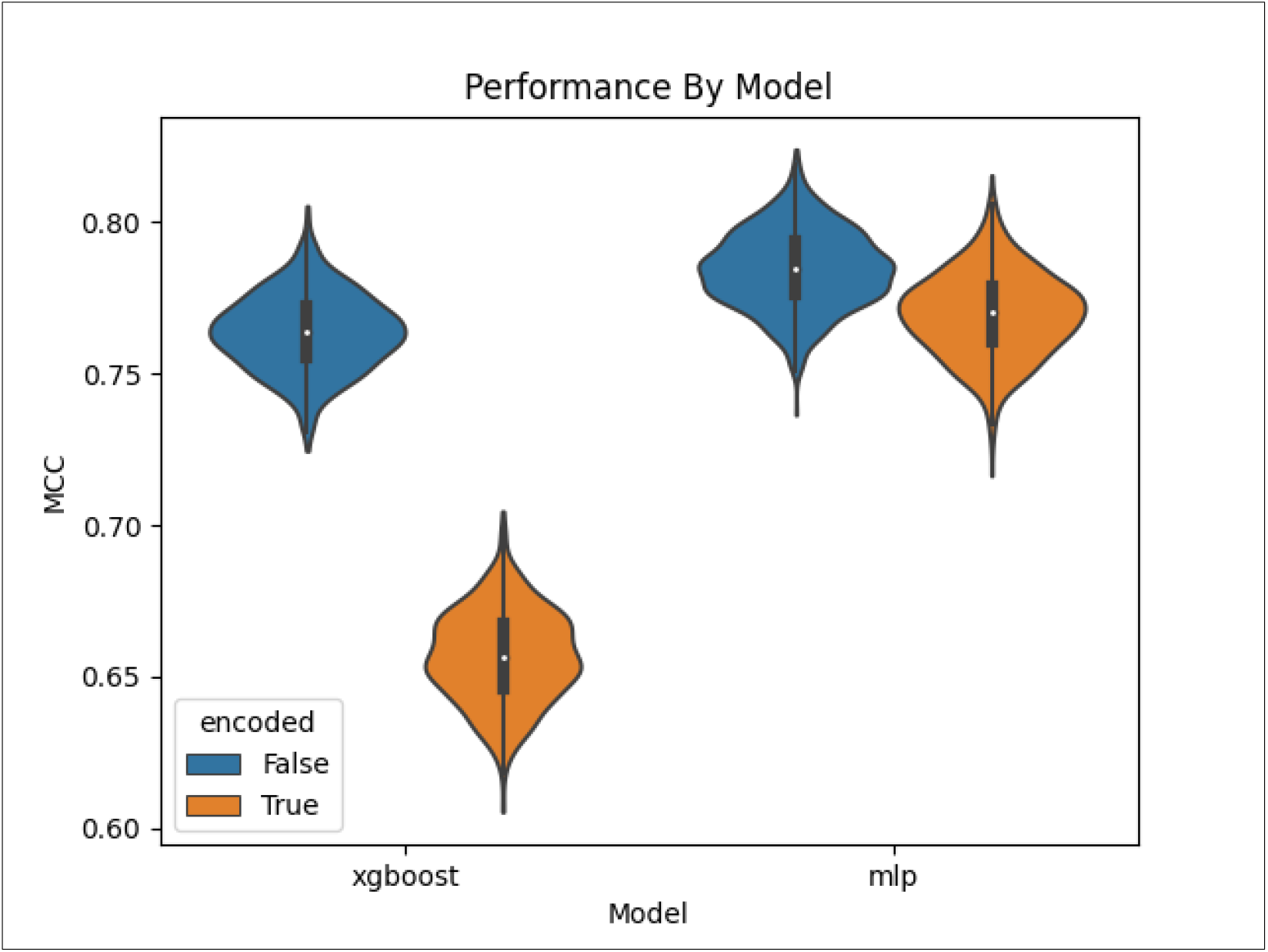
Overall performance by model.

To measure the computational resource usage of each model and for each feature set, we performed a subset of the CV iterations, this time while profiling the maximum RAM usage, the maximum GPU RAM, and the real compute time over 50 CV iterations. Table 5 details this information. For example, we see it took about 129 minutes for the MLP to complete the 50 iterations when training on the encoded data while taking about 90 minutes to do the same on the non-encoded data. The XGBoost took less time than the MLP when training on the same data. The non-encoded data moderately increased the GPU RAM and RAM utilization as compared to the encoded. The XGBoost required significantly more RAM and an order of magnitude more GPU RAM.

**Table 5.** Computational resource usage.

| Model | Computational resource |  |
| --- | --- | --- |
|  |  | Amount |
| MLP | RAM (gigabytes) | 2.507 |
|  | GPU RAM (gigabytes) | 0.848 |
|  | Compute time (minutes) | 89.706 |
| MLP with Encoding | RAM (gigabytes) | 1.764 |
|  | GPU RAM (gigabytes) | 0.42 |
|  | Compute time (minutes) | 129.130 |
| XGBoost | RAM (gigabytes) | 23.878 |
|  | GPU RAM (gigabytes) | 11.052 |
|  | Compute time (minutes) | 70.098 |
| XGBoost with Encoding | RAM (gigabytes) | 5.307 |
|  | GPU RAM (gigabytes) | 5.776 |
|  | Compute time (minutes) | 27.297 |

### 3.2. Feature Importance

Using the XGBoost model trained on the non-encoded feature set, we were able to compute the importance of each feature for each CV iteration. The softmax of the feature scores provided each feature importance relative to every other feature rather than the raw feature importance values. Taking the average relative feature importance across all CV iterations, every feature obtained an overall score indicating its feature importance. Those features scoring 0 were excluded from this analysis (Figure 7).

**Figure 7.**
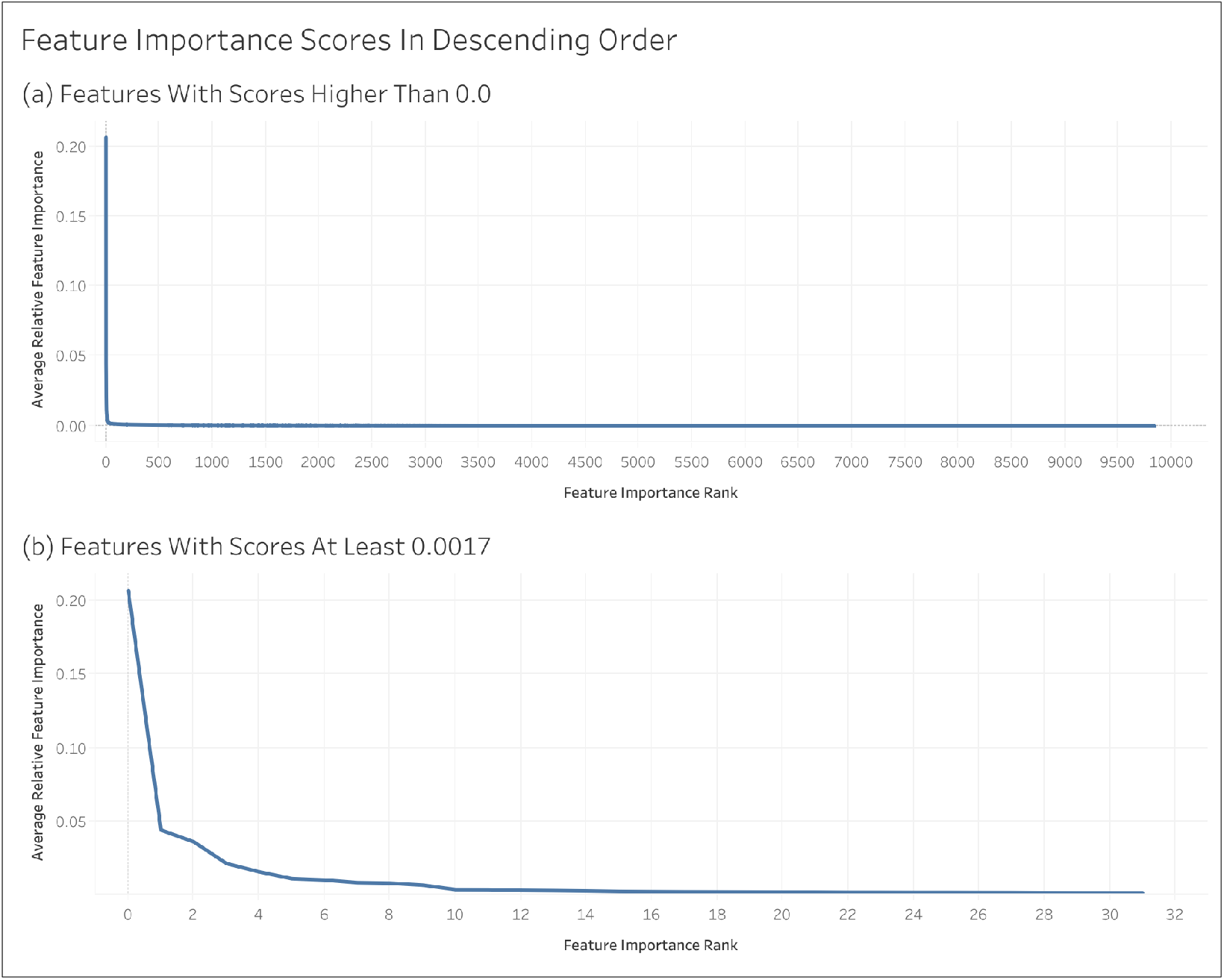
Feature importance scores in descending order.

Table 6 shows the atom colors [8] of the top 10 most important features, specifying whether the atom color corresponds to pathway features or to metabolite features. We see that most of the top 10 were pathway features. Associated pathways are defined by the atom color corresponding to a metabolite that exists in the pathway category. For example, the most important feature (rank 1) is associated with the ‘Biosynthesis of other secondary metabolites’, ‘Metabolism of terpenoids and polyketides’, and ‘Xenobiotics biodegradation and metabolism’ pathway categories. The rank of the feature is provided with 1 being the single most important feature down to the 10^th^ most important feature. If it’s a pathway feature, its pathway rank is the same as its rank. However, the corresponding metabolite feature (same atom color but representing metabolites instead of pathway categories) has its own rank. Both the pathway and metabolite rank are not always available either because the pathway feature corresponding to a metabolite feature may have been excluded from the feature set, having duplicate values as another feature. Or one of the features may have been a part of the feature set, not being a duplicate, but consistently scored 0 and wasn’t considered for feature importance. We see that for every pathway feature, the rank of its corresponding metabolite feature differs significantly..

**Table 6.**
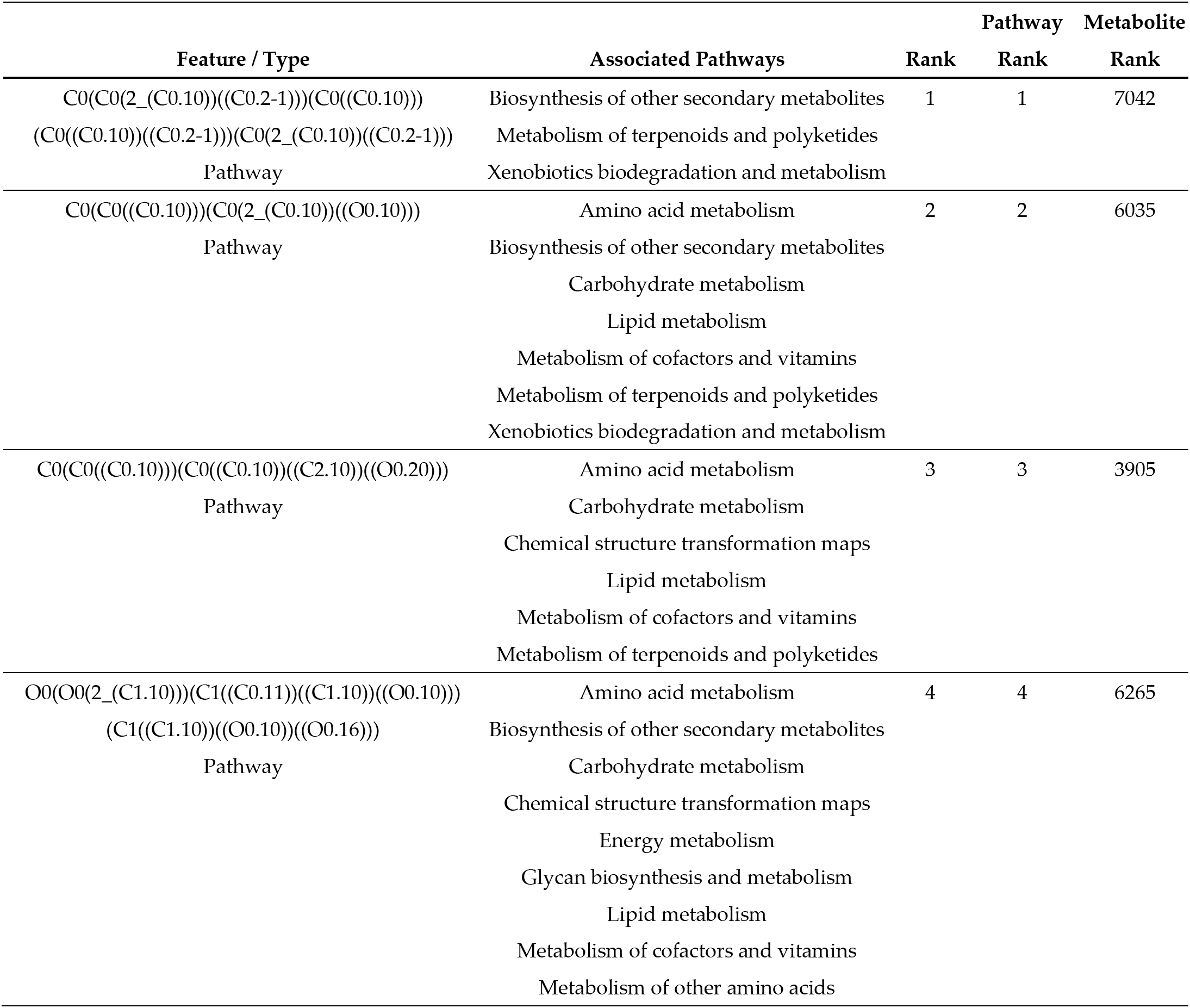
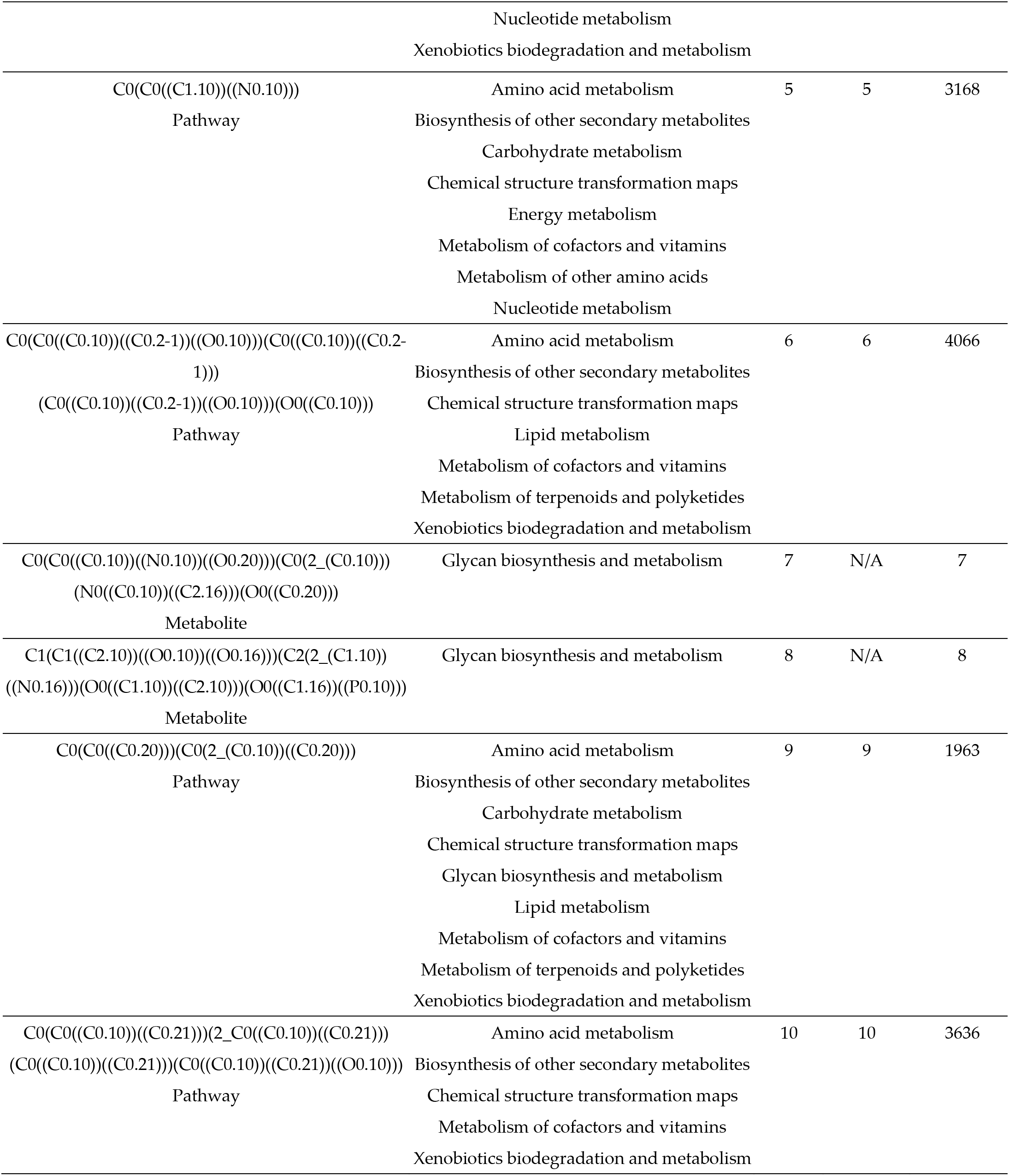
Top 10 most important features.

Figure 8 shows examples of compounds that contain the atom color corresponding to the top 10 most important features. The red highlighted portion contains the atoms and bonds corresponding to the atom color. We see the most important feature is a hydroxyl group connected to a methyl group by a ring structure.

**Figure 8.**
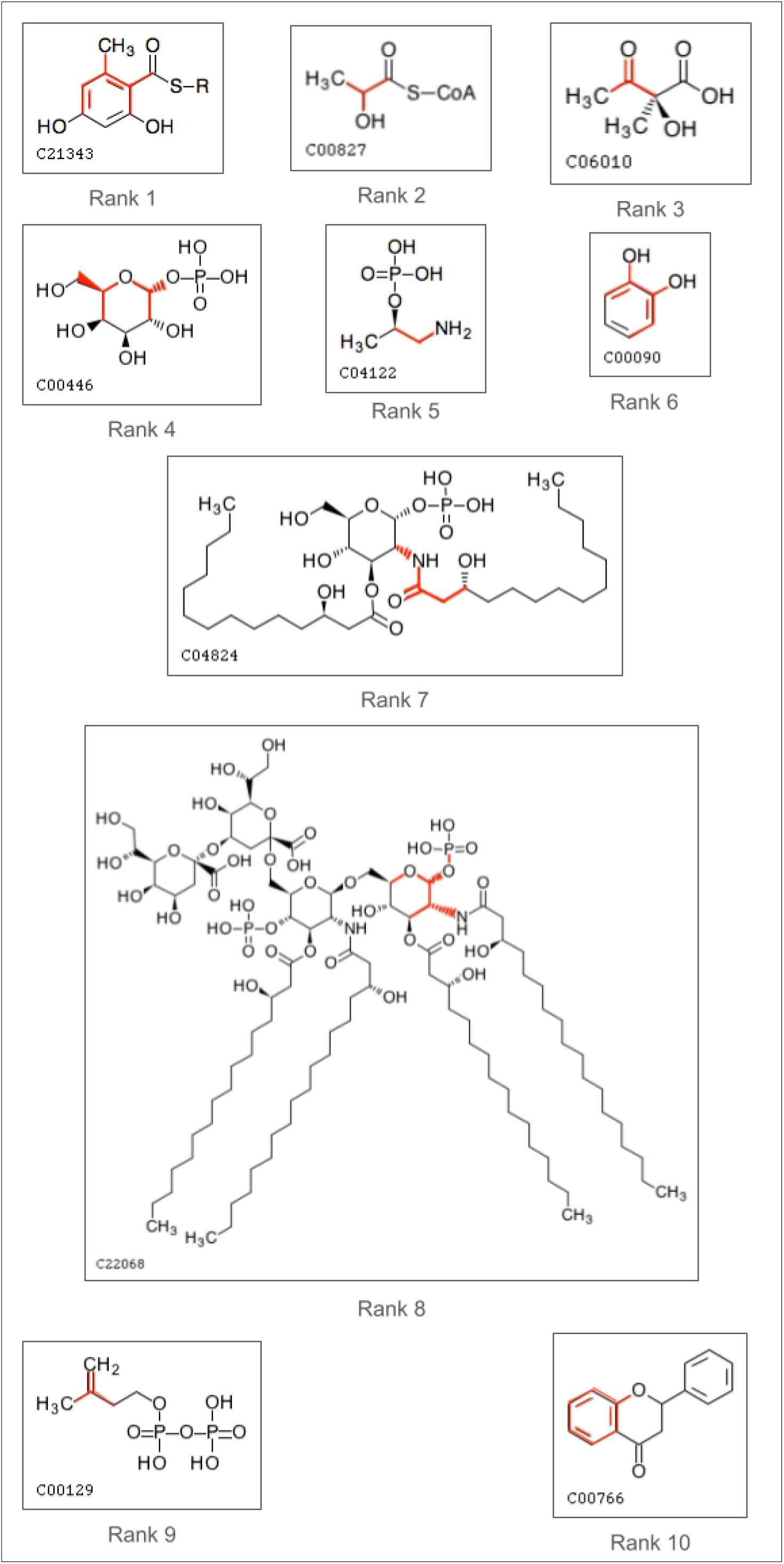
Molecular structures of top 10 features.

## 4. Discussion

Past publications interested in predicting pathway involvement of metabolites have trained separate binary classifiers for each specific pathway category in question, which has a number of shortcomings. In this work, we have overcome those shortcomings by training a single binary classifier that can predict the presence of a metabolite in a generically-described pathway category. The metabolite-pathway-pair technique simplifies the training, evaluation, and deployment process, reducing the computational resources required and enabling the possibility to predict involvement with arbitrarily defined pathway categories. From a literature search, we found a prior publication that utilized this metabolite-pathway-pair technique for designing a generic binary classifier for a different but similar classification problem predicting metabolite pathway involvement based on protein-protein interaction network data [32]. In our application predicting metabolite pathway involvement based on chemical structure, not only does the resulting single classifier perform comparably well to the combined performance of the separate classifiers, our best model has even exceeded the performance of the prior benchmark models (Table 4). In particular, our best model demonstrates greatly improved performance robustness as evidenced by the significant reduction of the standard deviation by an order of magnitude. We can confidently conclude that use of metabolite-pathway feature-pair entries is a superior method for predicting pathway involvement based on chemical structure derived features.

While XGBoost unsurprisingly performs relatively poorly when trained on the autoencoded features, we see that XGBoost trained on the non-encoded features performs comparably well with the MLP trained on the encoded features. The MLP when trained on the non-encoded features exceeds the performance of both as well as the combined performance of the prior benchmark models (Table 4). While the MLP requires a moderately larger amount of time to train and evaluate than the XGBoost, it requires only a fraction of the RAM and GPU RAM. And while the MLP trained on the non-encoded features requires a moderately larger amount of RAM and GPU RAM than that trained on the encoded features, the encoded variant actually requires significantly more time (Table 5). This is likely because while the non-encoded MLP takes longer to complete an individual epoch, the encoded MLP requires many more epochs to converge. Encoding the data saved RAM but even the non-encoded required less than 3 gigabytes of RAM and less than 1 gigabyte of GPU RAM, an extremely minor amount for modern HPC systems. With time being the more precious resource, we can confidently conclude that training the MLP on the non-encoded data is the best pipeline for this machine learning task and it appears that the autoencoder did not meaningfully save computational resources as initially expected.

When considering the importance of the atom-color features, there doesn’t appear to be any correlation between a pathway feature and its corresponding metabolite feature of the same atom color, and vice versa (Figure S1). Most of the top 10 features were pathway features, suggesting that the pathway information is often more important when predicting pathway involvement of a metabolite, as compared to information of the metabolite itself, at least from the perspective of XGBoost training.

Since the metabolite-pathway-pair approach is designed to scale to an arbitrarily high number of pathway categories, it paves the way for predicting more granular pathway categories as compared to the topmost 12 categories in the KEGG hierarchy. Beyond that, a model can be trained on pathway categories from KEGG combined with that from MetaCyc or any data source or any collection of arbitrary pathway categories, since the model is not tied to any particular category or restricted set of categories. This opens up the possibility to combine datasets derived from both KEGG and MetaCyc for future model development. Moreover, due to the multiplicative effect of the cross join when constructing metabolite pathway pairs, the dataset multiplies in size when adding more pathway categories. While this is typically a benefit for improving model performance, it may become impractical to train XGBoost models since they require the entire dataset fit into memory. However, neural networks train in mini-batches, making the MLP perhaps the more feasible model to use on future larger datasets, especially considering that the MLP requires less RAM and GPU RAM and outperforms XGBoost when given sufficient data. For smaller datasets, such as the one in this work, the XGBoost model may be preferred if analyzing feature importance is desired.

## Supplementary Materials

The following supporting information can be downloaded at:, Table 1: Hyperparameters, Figure S: Feature importance scores of pathway features compared to corresponding metabolite features, Table S2 - All Metrics And Scores.

## Author Contributions

Conceptualization, E.D.H and H.N.B.M; methodology, E.D.H.; software, E.D.H.; validation, E.D.H.; formal analysis, E.D.H.; investigation, E.D.H.; resources, H.N.B.M; data curation, E.D.H.; writing—original draft preparation, E.D.H.; writing—review and editing, H.N.B.M. and E.D.H.; visualization, E.D.H.; supervision, H.N.B.M.; project administration, H.N.B.M.; funding acquisition, H.N.B.M. All authors have read and agreed to the published version of the manuscript.”

## Funding

The research was funded by the National Science Foundation, grant number: 2020026 (PI Moseley), and by the National Institutes of Health, grant number: P42 ES007380 (University of Kentucky Super-fund Research Program Grant; PI Pennell). The content is solely the responsibility of the authors and does not necessarily represent the official views of the National Science Foundation nor the National Institute of Environmental Health Sciences.

## Institutional Review Board Statement

Not applicable.

## Informed Consent Statement

Not applicable.

## Data Availability Statement

The data and code for complete reproducibility of the results in this work are available via FigShare at: https://doi.org/10.6084/m9.figshare.25517695.v1

## Acknowledgments

We thank the University of Kentucky Center for Computational Sciences and Information Technology Services Research Computing for their support and use of the Lipscomb Compute Cluster and associated research computing resources.

## Conflicts of Interest

The authors declare no conflicts of interest.

---

**Table S1.**
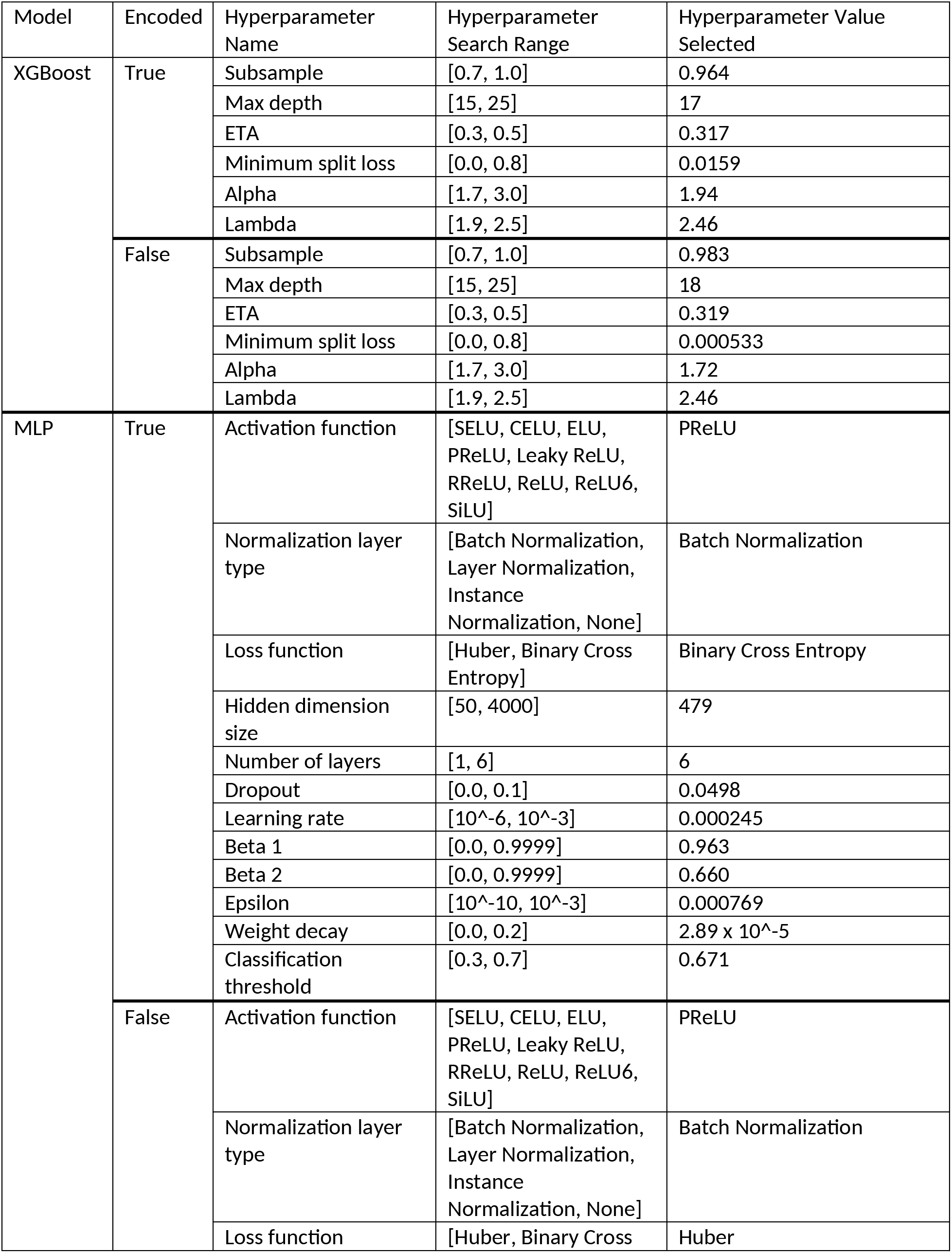
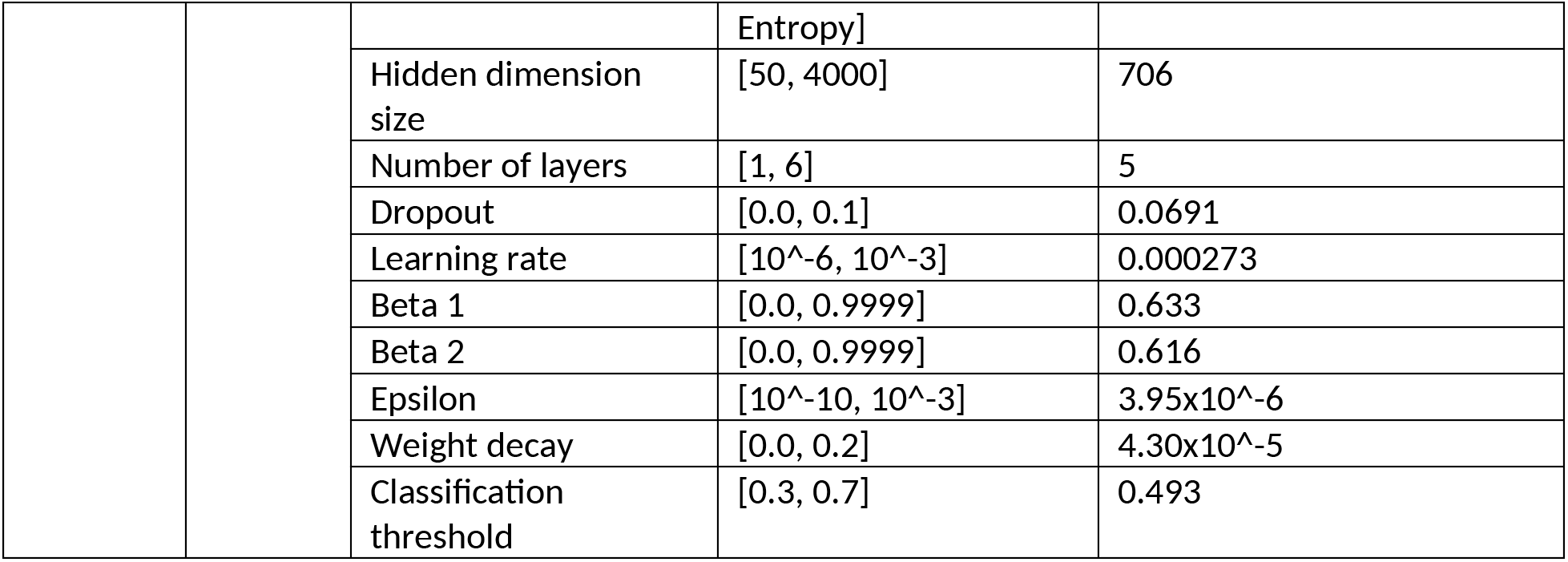
Hyperparameters.

**Figure S1.**
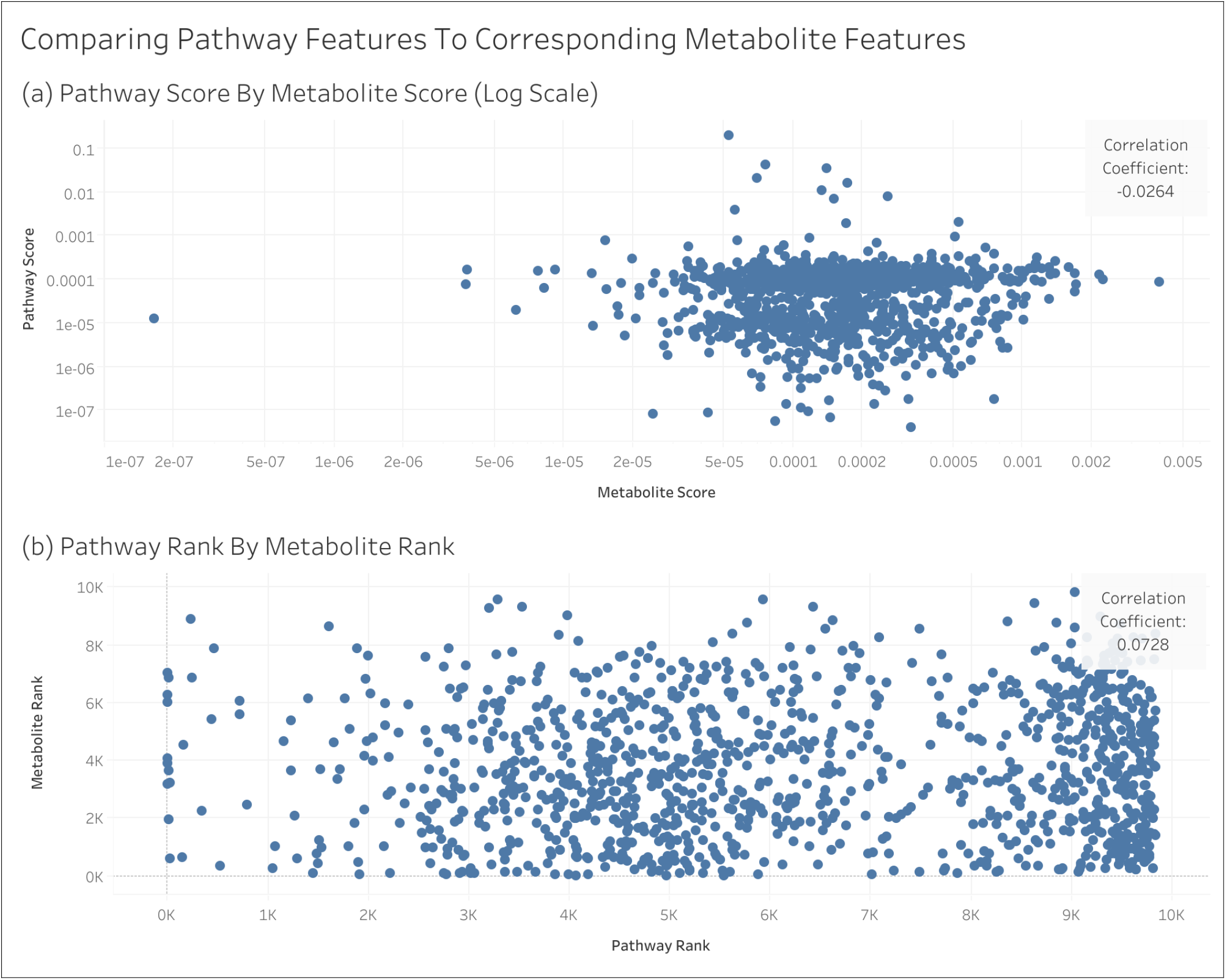
Feature importance scores of pathway features compared to corresponding metabolite features.

**Table S2.**
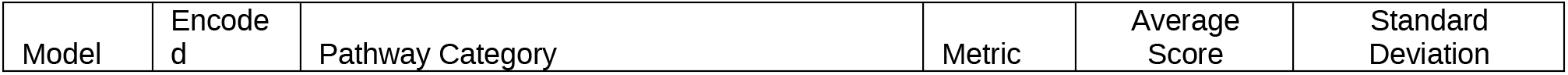
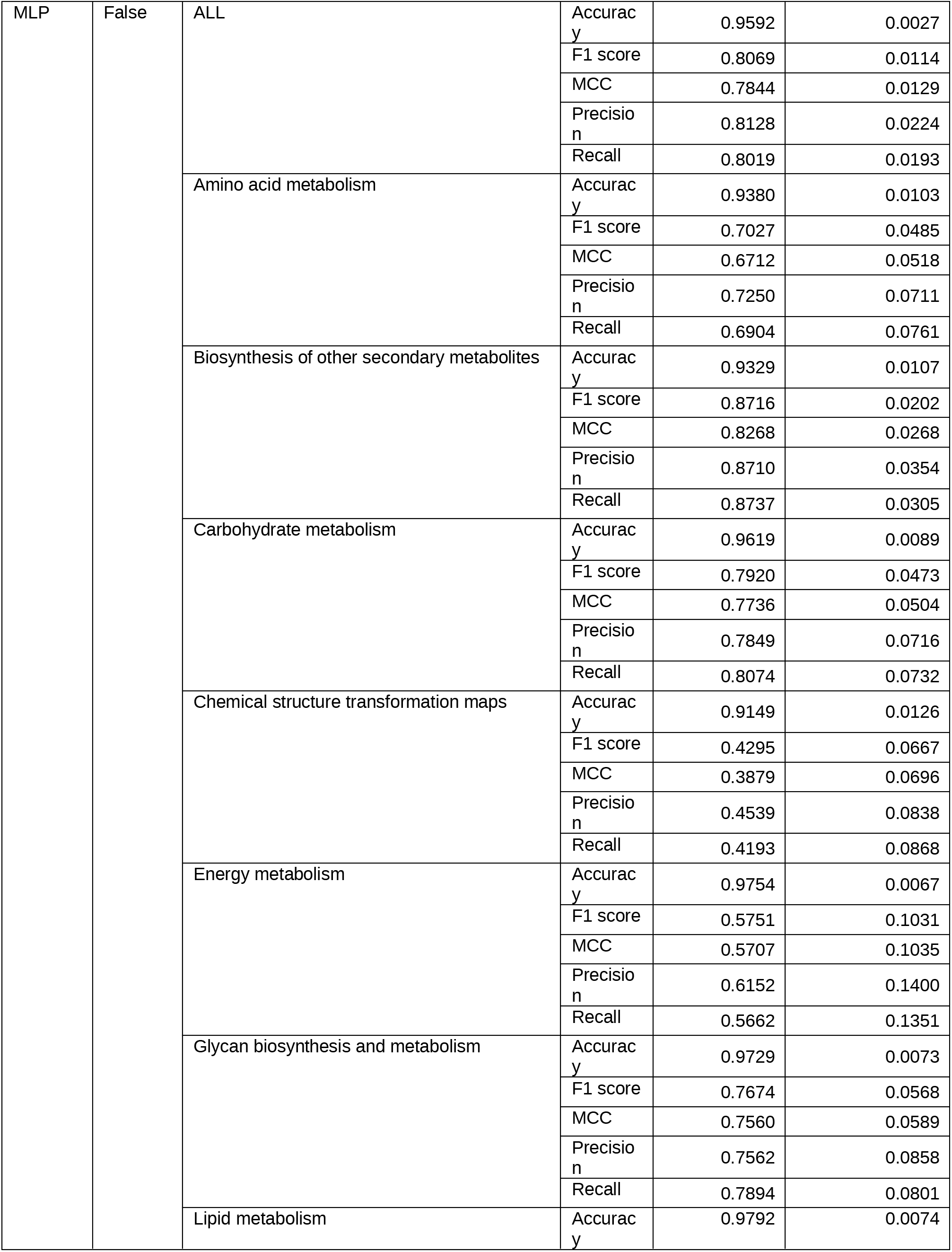
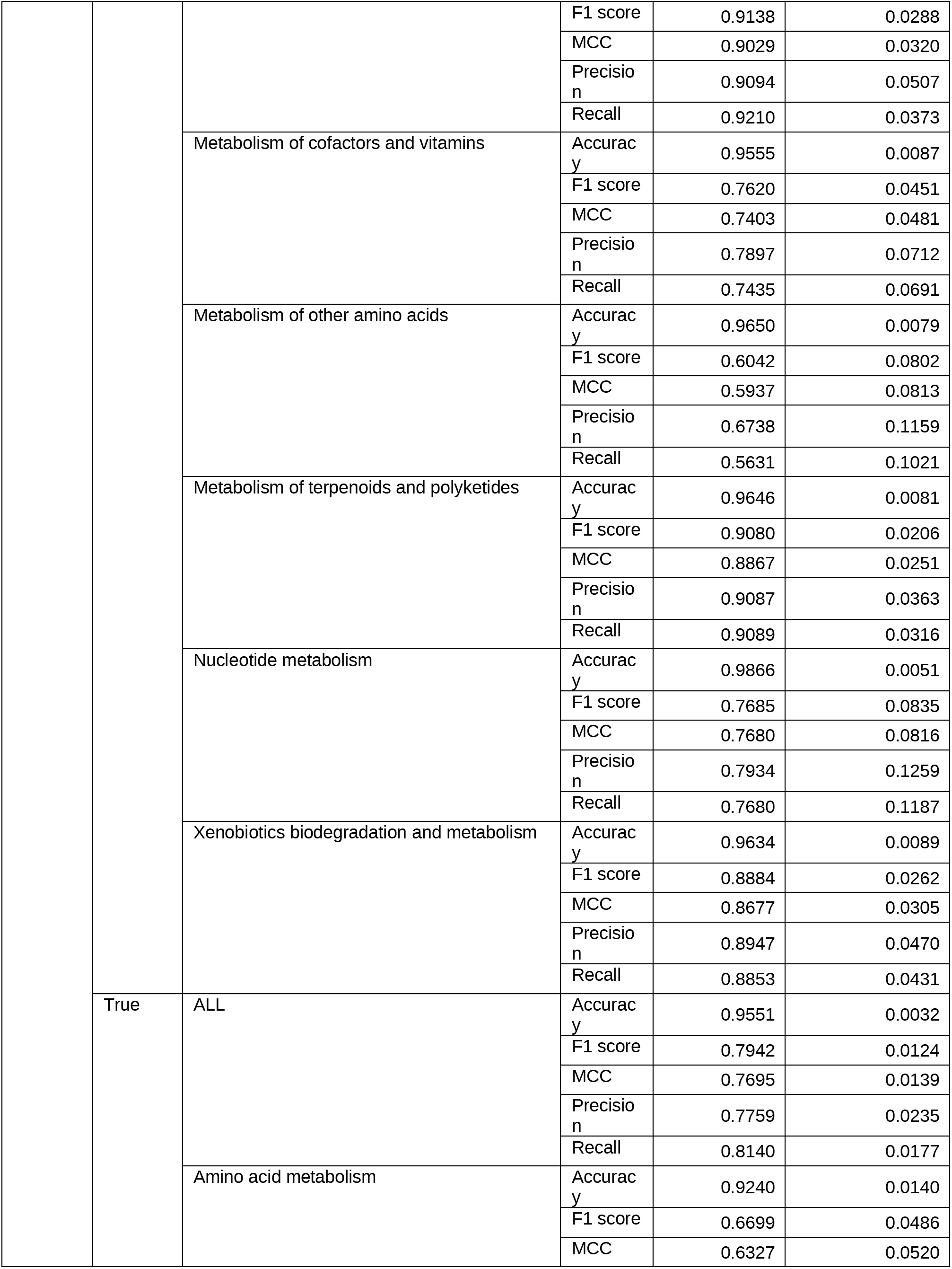
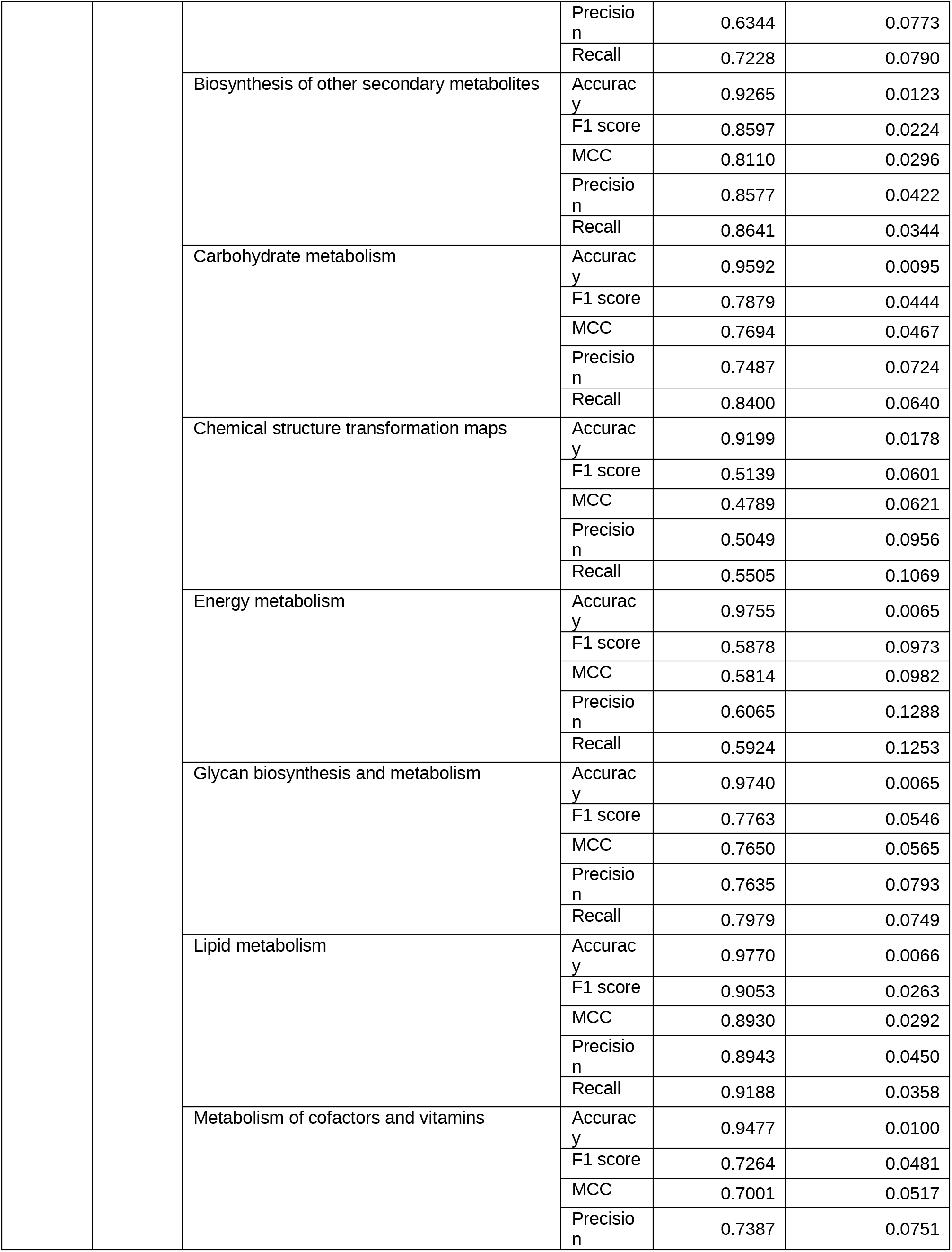
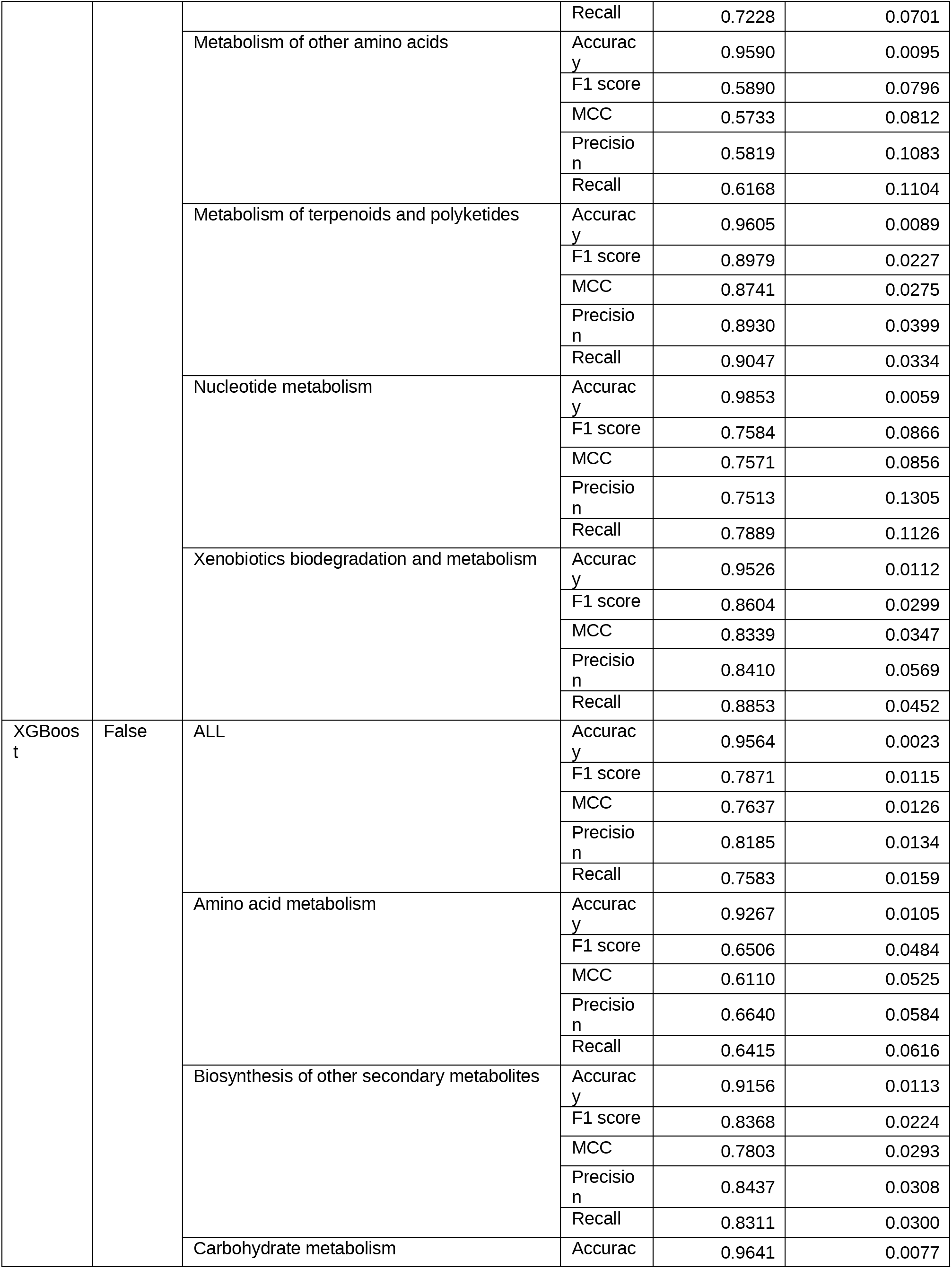
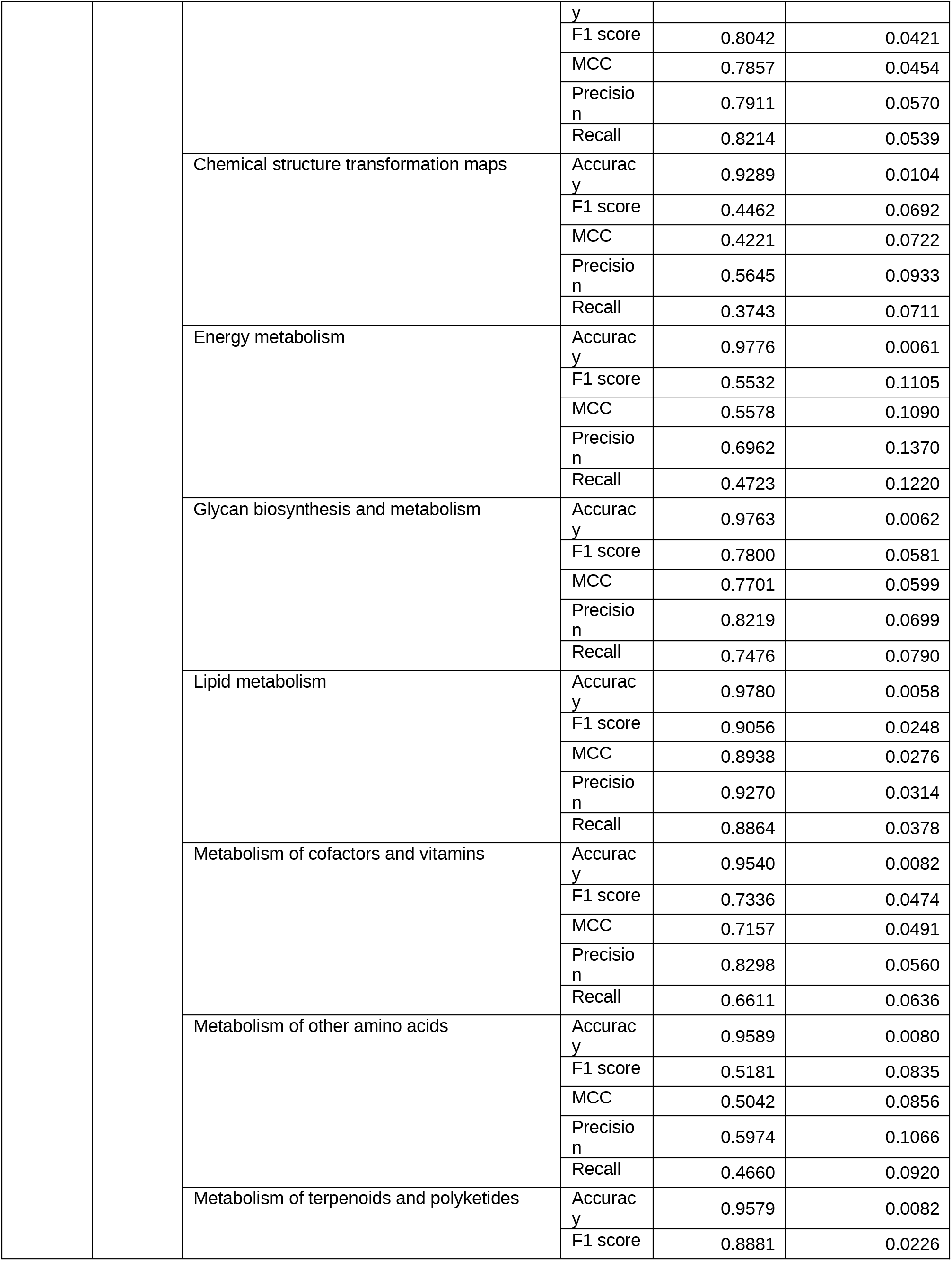
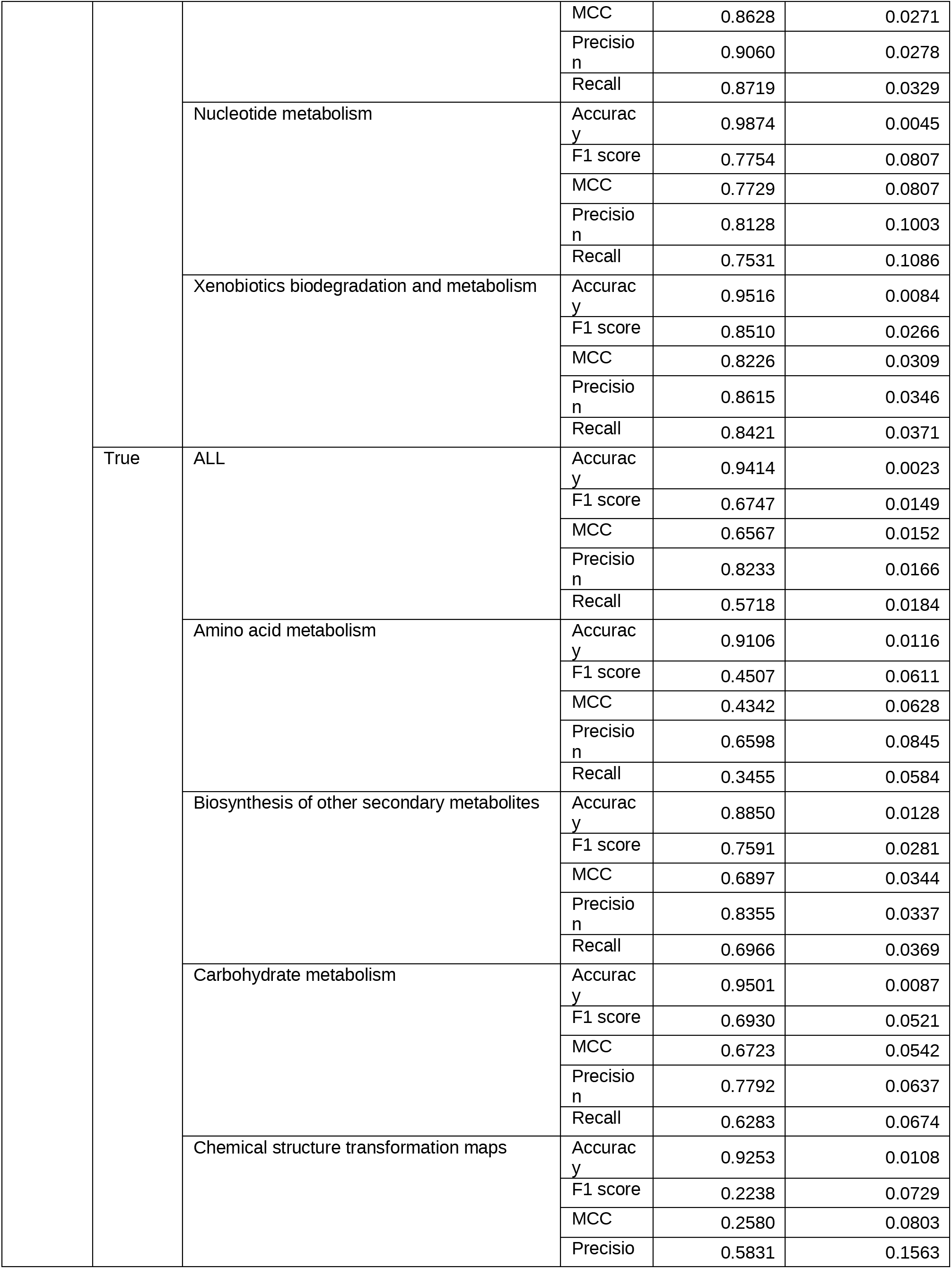
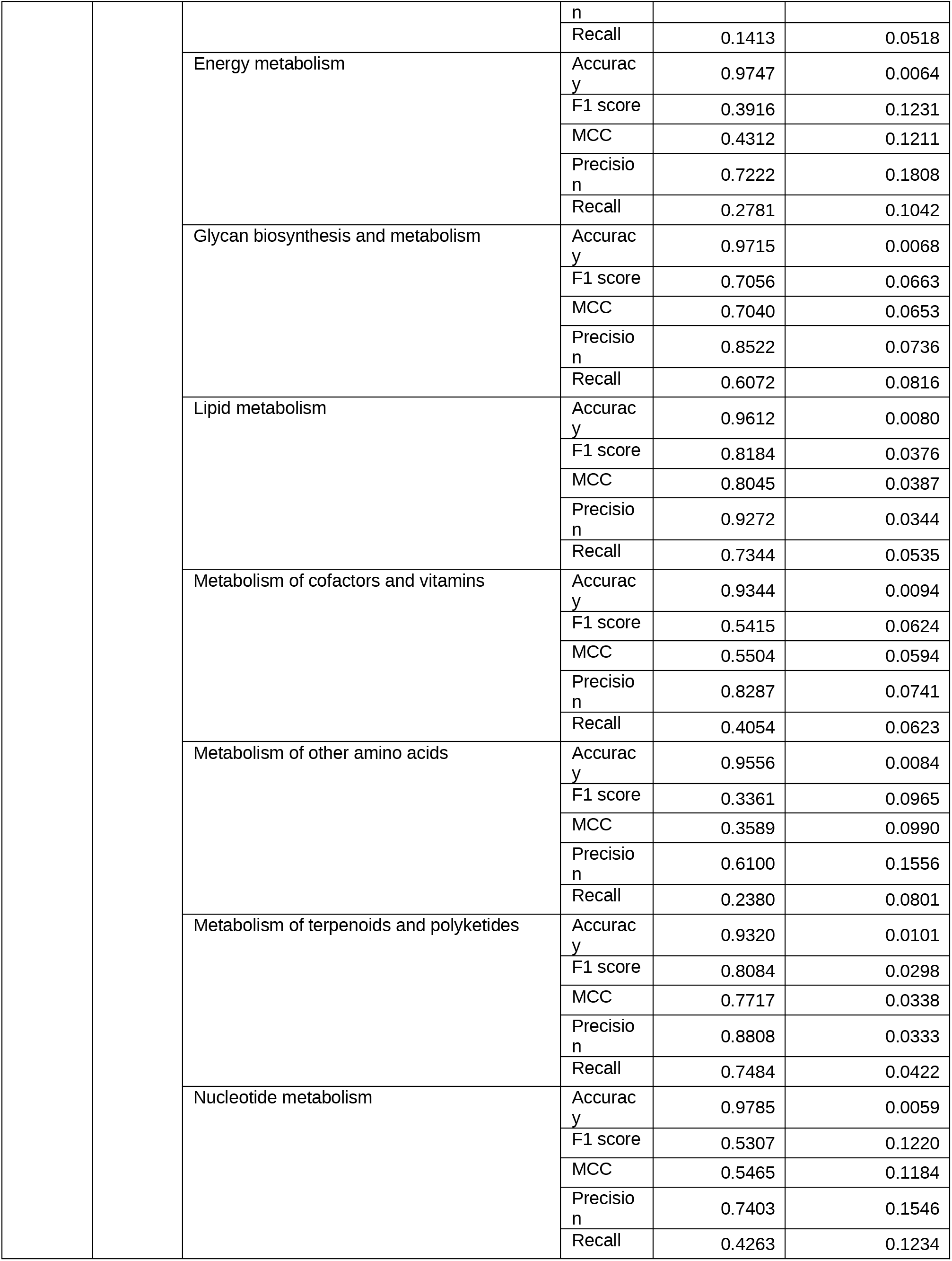
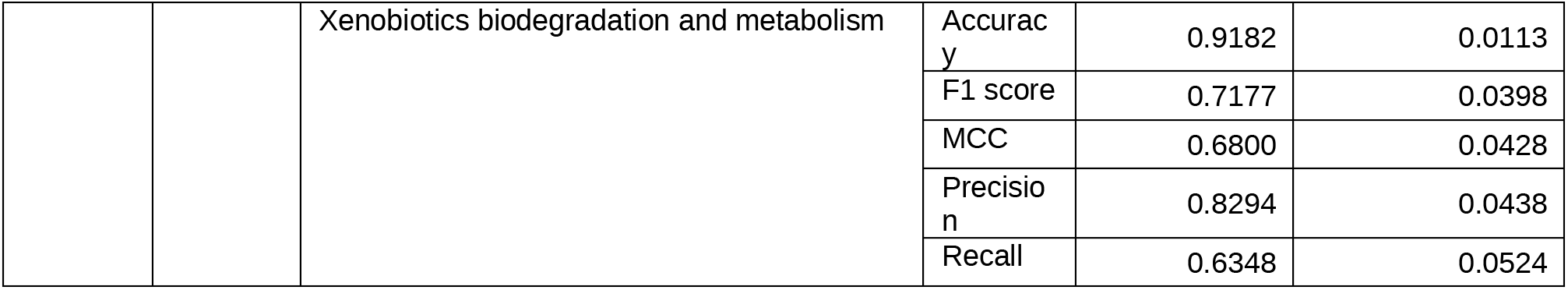
All Metrics And Scores.

## Notes

https://doi.org/10.6084/m9.figshare.25517695

### Competing Interest Statement

The authors have declared no competing interest.

